# A vapor exposure method for delivering heroin alters nociception, body temperature and spontaneous activity in female and male rats

**DOI:** 10.1101/2020.09.03.281857

**Authors:** Arnold Gutierrez, Kevin M. Creehan, Michael A. Taffe

## Abstract

**Background:** The ongoing crisis related to non-medical use of opioids makes it of continued importance to understand the risk factors for opioid addiction, the behavioral and neurobiological consequences of opioid exposure and to seek potential avenues for therapy. Pre-clinical rodent models have been critical to advancing understanding of opioid consequences for decades, but have been mostly limited to drug delivery by injection or by oral dosing. Inhalation, a significant route for many human users, has not been as well-established.

**Method:** We adapted an e-cigarette based exposure system, previously shown efficacious for delivery of other drugs to rats, to deliver heroin vapor. Effects *in vivo* were assessed in male and female Sprague-Dawley rats using a warm-water assay for anti-nociception and an implanted radiotelemetry system for evaluating changes in body temperature and spontaneous activity rate.

**Results:** Inhalation of vapor created by heroin 100 mg/mL in the propylene glycol (PG) vehicle significantly slowed tail-withdrawal from a 52°C water bath, bi-phasically altered activity, and increased temperature in male and female rats. Inhalation of heroin 50 mg/mL for 15 minutes produced significant effects, as the lower bound on efficacy, whereas inhalation of heroin 100 mg/mL for 30 minutes produced robust effects across all endpoints and groups.

**Conclusions:** This work shows that e-cigarette devices deliver psychoactive doses of heroin to rats, using concentrations of ∼50-100 mg/mL and inhalation durations of 15-30 minutes. This technique may be useful to assess the health consequences of inhaled heroin and other opioid drugs.

## 1. Introduction

The worldwide crisis related to the non-medical use of opioids continues, as reflected in high rates of deaths that are associated with prescription opioids, recreational drugs such as heroin and increasingly illicit supplies of fentanyl and analogs (CDC/NCHS, 2016; Wilson et al., 2020). The number of individuals seeking treatment for opioid dependence is high (SAMHSA, 2019). Improvement in understanding the risk factors for opioid addiction, as well as the behavioral and neurobiological consequences of opioid exposure, may enhance the discovery of new or improved avenues for therapy. Pre-clinical models in, e.g., laboratory rats, have been critical to advancing understanding but have been mostly limited to the examination of opioids delivered by parenteral injection, the insertion of subcutaneous pellets or pumps, or by oral dosing. Many humans use opioids by the inhalation route and indeed, one of the original public health crises related to non-medical use of opioids in the modern era was associated with the inhalation of vapor created by heating opium (Kane, 1881; Kritikos, 1960). The inhalation route might induce differences in the speed of brain entry, first pass metabolism, sequestration and release from non-brain tissues, etc. compared with these laboratory approaches. There is even some indication that inhalation use of heroin may come before a switch to injection use (Clatts et al., 2011; Cui et al., 2015; Rahimi-Movaghar et al., 2015; Syvertsen et al., 2017; Tyree et al., 2020), perhaps due to perceptions of safety or cultural factors. This may be true with respect to, e.g., disease transmission associated with intravenous injection practices but may involve other risks. For example, inhalation heroin users who develop leukoencephalopathy may be at increased risk for mortality compared with those who inject heroin (Alambyan et al., 2018; Cheng et al., 2019).

Prior studies have shown that the inhalation of heroin produces effects in animal models. Lichtman and colleagues reported anti-nociceptive effects of volatilized heroin and other opioids in mice using a heated glass pipe approach and a tail-withdrawal assay (Lichtman et al., 1996) and Carrol and colleagues demonstrated that monkeys would self-administer volatilized heroin (Mattox and Carroll, 1996; Mattox et al., 1997). There has, however, been no subsequent broad adoption of either of these techniques, possibly due to the difficulty of creating and maintaining what appear to be one-off apparatus/devices constructed by the respective lab groups. The recent advent of e-cigarette style devices (aka Electronic Nicotine Delivery Systems or ENDS) presents the possibility of delivering a wide range of drugs other than nicotine, including opioids, via vapor inhalation. These devices are widely available (Esterl, 2013; Farnham, 2013), offer significant translational appeal and can be easily adapted to use with laboratory rodents using commonly available sealed-top housing chambers, standard laboratory house vacuum, and simple plumbing. Human use of these devices for ingestion of cannabis extracts or cannabidiol (CBD) is becoming so popular (Jones et al., 2016; Kenne et al., 2017; Mammen et al., 2016; Morean et al., 2015; Morean et al., 2017; Poklis et al., 2019) that terming these devices e-cigarettes or ENDS is becoming a misnomer; Electronic Drug Delivery Systems (EDDS) is more accurate.

The primary goal of this study was to establish methods for delivering physiologically and behaviorally significant amounts of heroin to rats via vapor inhalation. To this end we adapted an e-cigarette based vaporization system we have previously shown is effective for the delivery of nicotine (Javadi-Paydar et al., 2019b), THC (Javadi-Paydar et al., 2018a; Nguyen et al., 2016b), cannabidiol (Javadi-Paydar et al., 2019a), methamphetamine (Nguyen et al., 2016a; Nguyen et al., 2017), 3,4-methylenedioxypyrovalerone (MDPV) (Nguyen et al., 2016a) and oxycodone (Nguyen et al., 2019) to rats. Other work has demonstrated anti-nociceptive effects of inhaled oxycodone in male rats (Nguyen et al., 2019) and a preliminary work demonstrates antinociceptive efficacy of inhaled heroin and methadone (Gutierrez et al., 2020) in male and female rats, and reinforcing effects of inhaled heroin in female rats. Nearly identical systems have been shown to deliver nicotine (Montanari et al., 2020), THC from cannabis extracts (Freels et al., 2020) and the potent synthetic opioids sufentanil and fentanyl (Moussawi et al., 2020; Vendruscolo et al., 2018), to laboratory rodents.

To validate the delivery of heroin by vapor inhalation, we used a warm water tail-withdrawal assay for assessing effects on nociception, and a radiotelemetry system capable of reporting body temperature and activity responses. The telemetry measures were selected based on prior evidence that parenteral injection of opioids causes hyperthermia and increased locomotor activity in both rats and mice. For example, intravenous morphine (6 mg/kg) increased the rectal temperature of anesthetized rats (El Bitar et al., 2016), as did intrathecal morphine (1-15 ug) in freely moving male Wistar rats measured with telemetry (Safrany-Fark et al., 2015). Both morphine and oxycodone produce hyperthermia when injected subcutaneously in rats (Bhalla et al., 2011). Solis and colleagues (2017) showed that intravenous fentanyl (40 ug/kg) initially decreased body temperature of rats in the first hour after administration, (heroin (400 ug/kg) did not), but each drug induced *hyper*thermia in the interval approximately 60-120 minutes after injection (Solis et al., 2017). Heroin has also been shown to increase the locomotor activity of rats when injected subcutaneously or intraperitoneally in doses of ∼0.25-1.0 mg/kg, although in some early studies a contrast of this described effect with vehicle injection was not made clear (Hoffman and Wise, 1993; Lamarque et al., 2001; Swerdlow and Koob, 1985; Swerdlow et al., 1985). Nevertheless, in other studies it was shown that heroin injection ∼0.25-0.5 mg/kg, s.c., significantly increases the locomotor activity of rats (Raleigh et al., 2013; Singh et al., 2005; Sorge and Stewart, 2006) in comparison with a vehicle injection.

A secondary goal of this study was to determine the parameters necessary for dose control / manipulation within an effective range. As we’ve shown in prior studies (Javadi-Paydar et al., 2019b; Javadi-Paydar et al., 2018a; Nguyen et al., 2016a; Nguyen et al., 2016b), this can be effected with changes in drug concentration in the vapor vehicle and the duration of exposure, thus these parameters were under investigation. The final goal was to determine if there are any substantive sex differences in the response to heroin inhalation, see Craft (Craft, 2008) for review, consistent with current suggestions for the inclusion of both male and female subjects in research where possible (Clayton and Collins, 2014; Shansky, 2020; Shansky and Woolley, 2016). While male and female rats acquire intravenous oxycodone self-administration at similar rates with 1-h access sessions (Mavrikaki et al., 2017), our recent study with 8-h access found that female rats self-administered more oxycodone in acquisition, but did not reach higher breakpoints in a PR dose-substitution (Nguyen et al., 2020b). A prior study reported no sex difference in the anti-nociceptive effect of heroin or oxycodone injected subcutaneously in Sprague-Dawley rats (Peckham and Traynor, 2006), thus no sex difference was predicted for this study.

## 2. Methods

### 2.1. Subjects

Female (N=7) and male (N=7) Sprague-Dawley rats (Harlan, Livermore, CA) rats were housed in humidity- and temperature-controlled (23±2 °C) vivaria on 12:12 hour light:dark cycles. Rats had *ad libitum* access to food and water in their home cages and all experiments were performed in the rats’ scotophase. Sample sizes were estimated based on results of prior studies investigating the effects of acute drug exposure on the telemetry endpoints (Aarde et al., 2017; Javadi-Paydar et al., 2018b; Nguyen et al., 2017; Nguyen et al., 2020a); group size was initially N=8 but two animals were initially lost due to surgical complications. All procedures were conducted under protocols approved by the Institutional Care and Use Committee of the University of California, San Diego.

### 2.2. Drugs

Heroin (diamorphine HCl) was administered by vapor exposure with doses described by the concentration in the propylene glycol (PG) vehicle (1, 50, 100 mg/mL) and duration of the exposure session (15, 30 minutes) as in prior studies (Javadi-Paydar et al., 2018a; Nguyen et al., 2016b). Heroin (0.25, 0.5, 1.0 mg/kg) was dissolved in physiological saline for subcutaneous injection. The heroin was provided by the U.S. National Institute on Drug Abuse Drug Supply Program.

### 2.3. Exposure Apparatus

Vapor was delivered into sealed vapor exposure chambers (152 mm W X 178 mm H X 330 mm L; La Jolla Alcohol Research, Inc, La Jolla, CA, USA) through the use of e-vape controllers (Model SSV-3 and SVS-200; 58 watts, 0.24-0.26 ohms, 3.95-4.3 volts, ∼214 °F; La Jolla Alcohol Research, Inc, La Jolla, CA, USA) to trigger Smok Baby Beast Brother TFV8 sub-ohm tanks. Tanks were equipped with V8 X-Baby M2 0.25 ohm coils. Timed activation of e-vape controllers to deliver the scheduled drug vapor (one 6 sec puff every 5 minutes) was done using MedPC IV software (Med Associates, St. Albans, VT USA). The chamber air was vacuum-controlled by a chamber exhaust valve (i.e., a “pull” system) to flow room ambient air through an intake valve at ∼1 L per minute. Vacuum was turned on 30 seconds prior to the vapor triggering to clear any vapor from the prior puff. The vacuum remained on for 10 seconds (i.e. during, and for 4 seconds after, each puff), to ensure that vapor filled the chamber.

### 2.4. Radiotelemetry

The rats were implanted with sterile radiotelemetry transmitters (TA11TA-F20; Data Sciences International, St Paul, MN) in the abdominal cavity as previously described (Taffe et al., 2015; Wright et al., 2012) on Post-Natal Day 31. For studies, animals were evaluated in clean standard plastic homecages (∼1 cm layer of sani-chip bedding) in a dark testing room, separate from the vivarium, during the (vivarium) dark cycle. Radiotelemetry transmissions were collected via telemetry receiver plates (Data Sciences International, St Paul, MN; RPC-1 or RMC-1) placed under the cages as described in prior investigations (Javadi-Paydar et al., 2018b; Nguyen et al., 2017). Test sessions for exposure studies started with a 15-minute interval to ensure data collection, then a 15-minute interval for baseline temperature and activity values. The rats were then moved to a separate room for the vapor exposure sessions and then returned to the recording chambers for up to 300 minutes after the start of vapor exposure. These experiments were conducted over the adult age interval of PND 204-235. The subjects had participated in similar telemetry recording experiments following vapor and injected exposure to doses of nicotine and THC from PND 50 to PND 190 with active dosing conducted no more frequently than two times per week, in studies not previously reported.

### 2.5 Experiments

#### 2.5.1 Experiment 1: Temperature and Activity

The rats were recorded for a baseline interval and were then exposed to vapor (Heroin 0, 1, 50 mg/mL in the PG) for the designated interval (15 or 30 minutes) in exposure subgroups of 2-3 rats of the same sex per chamber. The initial dose range was determined by pilot studies and adaptation from self-administration studies published at present only in pre-print form (Gutierrez et al., 2020). The vapor conditions were evaluated in a counterbalanced order by subgroup, with exposure duration alternating each session by sex. Sessions were conducted up to twice per week, i.e., no more frequently than every 3-4 days. Following this, the effects of Heroin 100 mg/mL for 15 vs 30 minutes, alternating by sex, were assessed on two different days, again at a 3-4 day interval.

#### 2.5.2 Experiment 2: Anti-nociception

A tail-withdrawal assay for nociception was conducted immediately after the vapor sessions, i.e. during Experiment 1, before the rats were returned to their individual recording chambers for telemetry assessment. The latency to withdraw the tail from insertion (3-4 cm) into a 52°C water bath was recorded by stopwatch, using methods as previously described (Gutierrez et al., 2020; Javadi-Paydar et al., 2018a; Nguyen et al., 2019). A 15 sec cutoff was used as the maximal latency for this assay to avoid any potential for tissue damage.

#### 2.5.1 Experiment 3: Subcutaneous injection

The rats were next recorded for a baseline interval and were injected with saline or Heroin (0.25, 0.5, 1.0 mg/kg, s.c.) in a counter-balanced order. Following injection the rats were immediately replaced in the recording chambers. Sessions were conducted up to twice per week, i.e., no more frequently than every 3-4 days. One male rats’ transmitter stopped functioning correctly during this experiment thus N=6 for this group.

### 2.6. Data Analysis

The telemeterized body temperature and activity rate (counts per minute) were collected on a 5-minute schedule in telemetry studies but are expressed as the average of six sequential measurements (referred to as “30-minute” averages) for primary analysis. The time courses for data collection are expressed relative to the initiation of vapor exposure and times in the figures refer to the end of the interval (e.g. “90 minutes” reflects the data collected from 65 to 90 minutes, inclusively. Due to transfer time after vapor sessions, the “60 minutes” interval reflects the average of collections from ∼40-60 minutes). Any missing temperature values were interpolated from the values before and after the lost time point. Activity rate values were not interpolated because 5-minute to 5-minute values can change dramatically, thus there is no justification for interpolating. Data for the first time-point after the start was inadvertently missing for two female rats in the 30-minute PG condition. The values for the second timepoint were used in place of the missing values. These ended up being very similar to the respective values recorded for this timepoint for the 15-minute PG condition. The telemetry data (temperature, activity rate) were generally analyzed with two-way Analysis of Variance (ANOVA) including repeated measures factors for the Vapor condition and the Time after vapor initiation. A mixed-effects model was used instead of ANOVA wherever there were missing datapoints. The nociception data were analyzed by three-way ANOVA including repeated-measures factors for Duration of exposure and Vapor condition and a between-groups factor of Sex. Any significant main effects were followed with post-hoc analysis using Tukey (multi-level factors) or Sidak (two-level factors) procedures. All analysis used Prism 8 for Windows (v. 8.4.3; GraphPad Software, Inc, San Diego CA).

## 3. Results

### 3.1 Experiment 1 (Telemetry)

#### 3.1.1 15-minute inhalation

##### Female Rats

The body temperature of the female rats (**Figure 1A**) was significantly elevated by the inhalation of Heroin 50 or 100 mg/mL as reflected in a significant effect of Time after vapor initiation [F (9, 54) = 13.97; P<0.0001] and of the interaction of Time with Vape Condition [F (27, 162) = 5.91; P<0.0001] on temperature. The post-hoc test confirmed that temperature was significantly higher after inhalation of Heroin 50 mg/mL compared with the PG condition (60-150 minutes after the start of inhalation) and Heroin 1 mg/mL (60-120 minutes) and likewise higher after inhalation of Heroin 100 mg/mL from 90-150 minutes compared with both the PG and Heroin 1 mg/mL inhalation conditions. Relatedly, temperature was lower than the baseline after inhalation of PG (120-210 minutes after the start of inhalation) or Heroin 1 mg/mL (90-180; 300 minutes) inhalation conditions, but was not significantly changed from baseline in the Heroin 50 mg/mL condition until 180 and 270 minutes after the start of inhalation and from 180-300 minutes after the start of Heroin 100 mg/mL inhalation.

**Figure 1:**
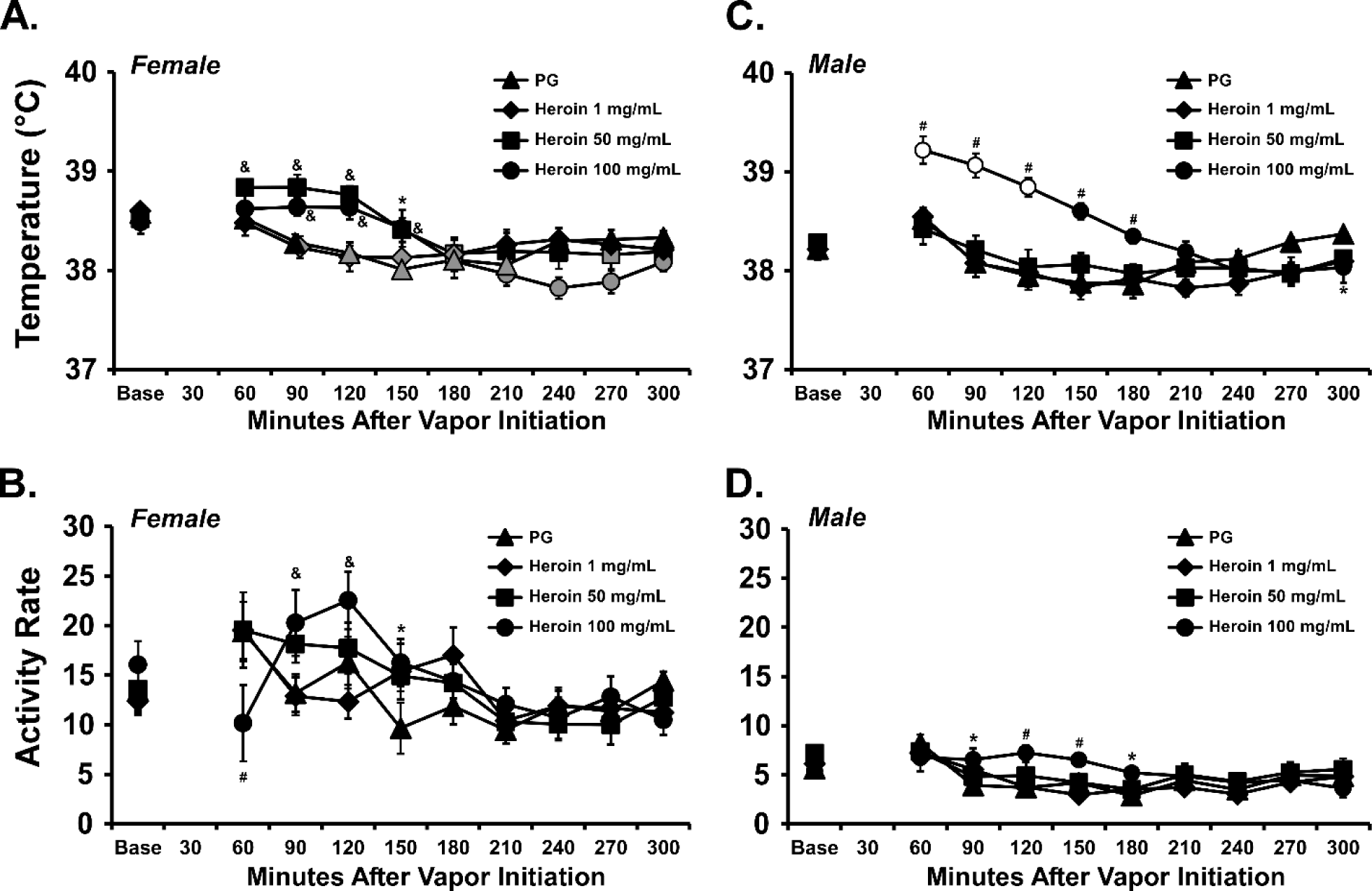
Mean (N=7 per group; ±SEM) body temperature (A, C) and activity rate (B, D) for female (A, B) and male (C,D) Sprague-Dawley rats after inhalation of vapor from the propylene glycol vehicle (PG), or Heroin (1, 50, 100 mg/mL in the PG) for 15 minutes. Shaded symbols indicate a significant difference from the baseline (Base) value, open symbols indicate a significant difference from the baseline and the PG condition at the respective time-point. A significant difference from the PG condition at a given time is indicated with *, a difference from PG and Heroin 1 mg/mL with &, and a difference from all other treatment conditions with #.

There was also a significant effect of Time [F (9, 54) = 7.42; P<0.0001] and of the interaction of Time with Vape Condition [F (27, 162) = 2.89; P<0.0001] on the activity of the female rats (**Figure 1B**). The post-hoc test confirmed that female rat activity was lower 60 minutes after the start of inhalation of Heroin 100 mg/mL compared with all other conditions and thereafter higher compared with PG (90-150 minutes) or Heroin 1 mg/mL (90-120 minutes) inhalation.

##### Male rats

The body temperature of male rats (**Figure 1C**) was significantly altered by Time after vapor initiation [F (9, 54) = 26.60; P<0.0001], by Vapor Condition [F (3, 18) = 7.07; P<0.005] and of the interaction of Time with Vapor Condition [F (27, 162) = 6.41; P<0.0001]. The post-hoc test confirmed that temperature was higher after inhalation of Heroin 100 mg/mL compared with all three other inhalation conditions from 60-180 minutes after the start of inhalation and lower compared with PG inhalation at the 300-minute time-point. Body temperature differed significantly from the baseline after inhalation of Heroin 100 mg/mL (60-120 minutes) but not in the other inhalation conditions. There was a significant effect of Time [F (9, 54) = 13.86; P<0.0001] and of the interaction of Time with Vape Condition [F (27, 162) = 2.37; P<0.001] on the activity of the male rats (**Figure 1D**). The post-hoc test confirmed that activity was greater after inhalation of Heroin 100 mg/mL compared with inhalation of PG (90-180 minutes after the start of inhalation) and compared with the other two heroin conditions (120-150 minutes).

#### 3.1.2 30-minute inhalation

##### Female Rats

The body temperature of the female rats (**Figure 2A**) was affected by Heroin inhalation for 30 minutes, as confirmed by a significant effect of Time after vapor initiation [F (9, 54) = 6.98; P<0.0001] and of the interaction of Time with Vape Condition [F (27, 162) = 3.19; P<0.0001] on temperature in the ANOVA. The post-hoc test confirmed that temperature was lower 60-90 minutes after the start of inhalation of Heroin 100 mg/mL compared with the PG (and 60 and 90 minutes after the start of the Heroin 50 mg/mL and Heroin 1 mg/mL conditions, respectively). Temperature was also lower than the baseline 60 minutes after initiation of Heroin 1 or 100 mg/mL. Temperature was then *higher* than in the PG condition after inhalation of Heroin 50 or 100 mg/mL, 120-150 minutes after the start of inhalation. The biphasic nature of the temperature response in the female rats is better illustrated by the 5-minute sample resolution for individuals presented in **Figure 3**.

**Figure 2:**
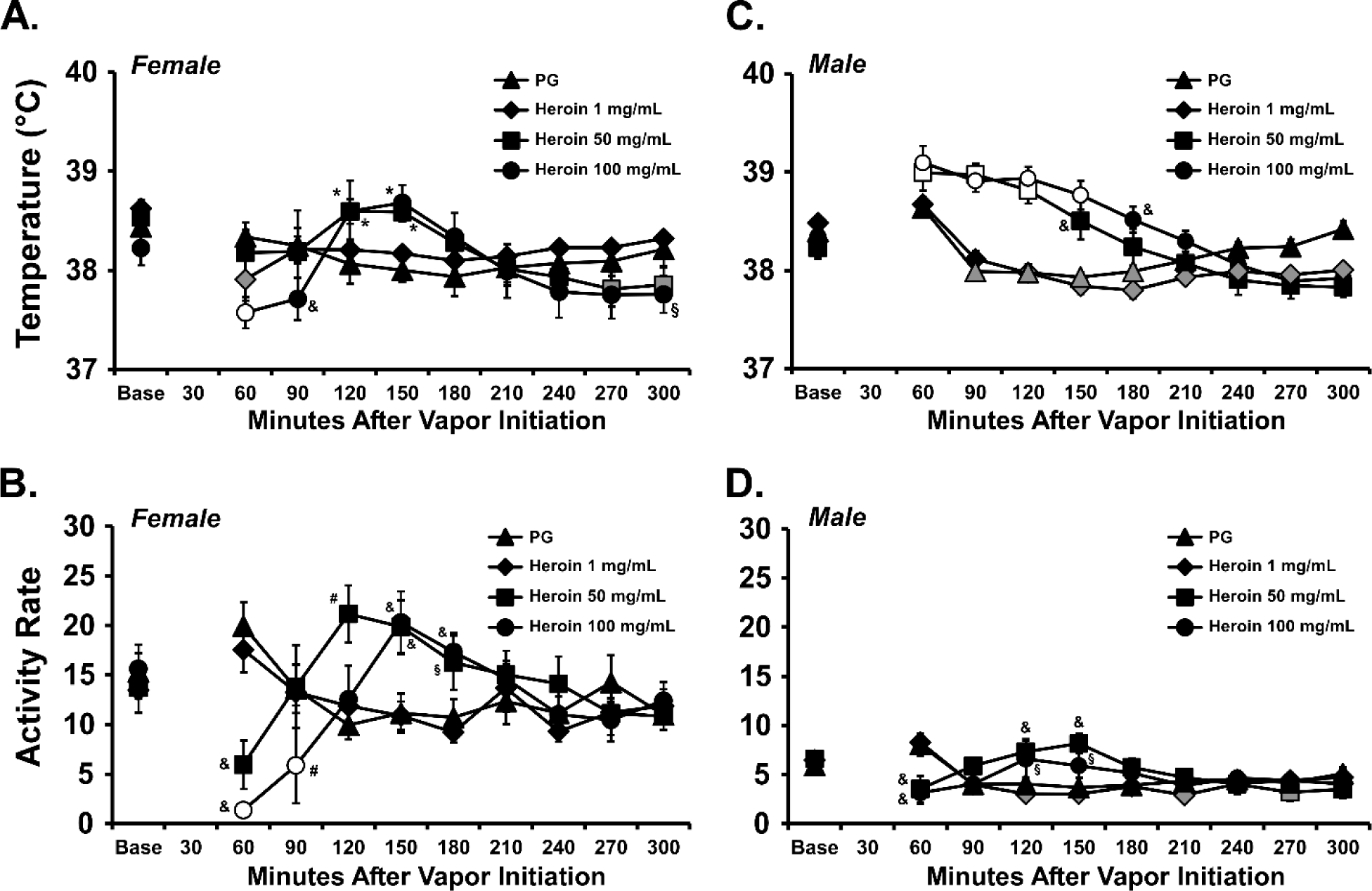
Mean (N=7 per group; ±SEM) body temperature (A, C) and activity rate (B, D) for female (A, B) and male (C,D) Sprague-Dawley rats after inhalation of vapor from the propylene glycol vehicle (PG), or Heroin (1, 50, 100 mg/mL in the PG) for 30 minutes. Shaded symbols indicate a significant difference from the baseline (Base) value, open symbols indicate a significant difference from the baseline and the PG condition at the respective time-point. A significant difference from the PG condition at a given time is indicated with *, a difference from Heroin 1 mg/mL with §, a difference from PG and Heroin 1 mg/mL with &, and a difference from all other treatment conditions with #.

**Figure 3:**
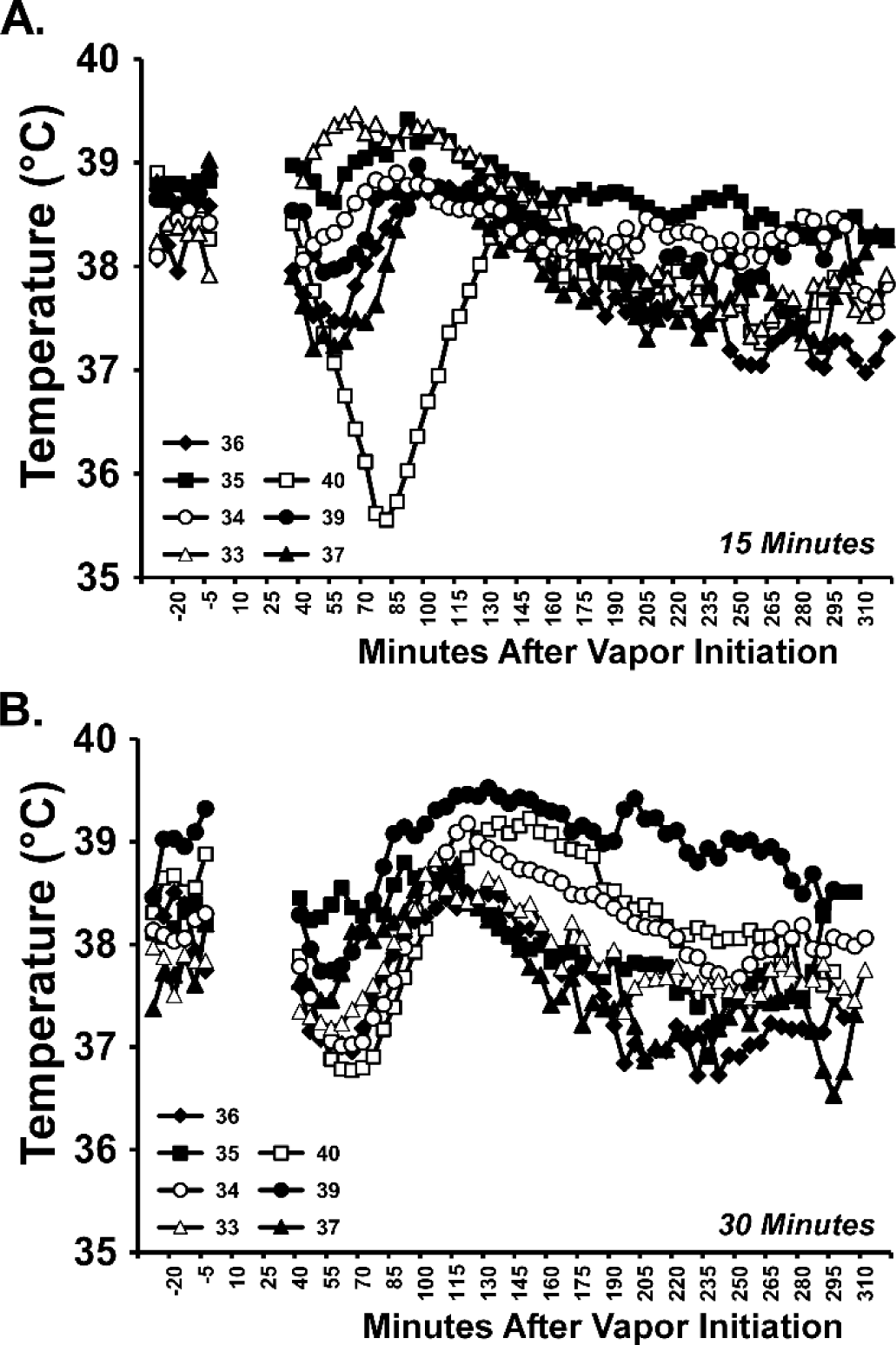
Individual body temperature responses in the female group after inhalation of Heroin 50 and 100 mg/mL for A) 15 minutes and B) 30 minutes illustrates the consistently biphasic nature of the temperature response.

The activity of female rats (**Figure 2B**) was also altered by Heroin inhalation and the ANOVA confirmed significant effects of Time after vapor initiation [F (9, 54) = 3.24; P<0.005] and of the interaction of Time with Vape Condition [F (27, 162) = 6.09; P<0.0001] on activity rate. Activity was suppressed after Heroin 50 mg/mL (60 minutes after the initiation of vapor) or 100 mg/mL (60-90 minutes) vapor exposure compared with the PG and Heroin 1 mg/mL conditions. Activity was also significantly lower than the baseline 60-90 minutes after the start of 100 mg/mL inhalation, and lower compared with the 50 mg/mL condition in the 90-minute time-point. Activity rates were thereafter significantly higher compared with all other conditions after the inhalation of Heroin 50 mg/mL (120 minutes), and higher compared with the PG or Heroin 1 mg/mL after Heroin 50 mg/mL (150 minutes) or 100 mg/mL (150-180 minutes) conditions. Activity differed significantly between Heroin 50 mg/mL and Heroin 1 mg/mL inhalation 180 minutes after the start of inhalation as well.

##### Male Rats

The body temperature of the male rats (**Figure 2C**) was significantly affected by Heroin inhalation for 30 minutes, and the ANOVA confirmed significant effects of Time after vapor initiation [F (9, 54) = 43.43; P<0.0001], of Vapor Condition [F (3, 18) = 5.16; P<0.01], and of the interaction of Time with Vapor Condition [F (27, 162) = 11.06; P<0.0001] on temperature. The post-hoc test confirmed that body temperature of the male rats was elevated, compared with the respective time after PG or Heroin 1 mg/mL inhalation, after the Heroin 50 mg/mL (60-150 minutes after the start of inhalation) or Heroin 100 mg/mL (60-180 minutes) inhalation (**Figure 2B**). Similarly, temperature was *higher* than the baseline in the Heroin 50 mg/mL (60-120 minutes after the start of inhalation) and Heroin 100 mg/mL (60-150 minutes) whereas it was *lower* than the baseline for each of the PG (90-180 minutes) and Heroin 1 mg/mL (120-300 minutes) inhalation conditions.

The activity of male rats (**Figure 2D**) was much lower than that of the female rats, but was also altered by Heroin inhalation and the ANOVA confirmed significant effects of Time after vapor initiation [F (9, 54) = 5.69; P<0.0001] and of the interaction of Time with Vape Condition [F (27, 162) = 4.06; P<0.0001] on activity rate. As with the female rats, the activity of the males was suppressed after Heroin 50 or 100 mg/mL (60 minutes after the initiation of vapor) vapor inhalation compared with the PG and Heroin 1 mg/mL conditions. Activity rates were thereafter significantly higher compared with PG inhalation in the Heroin 50 mg/mL (120-150 minutes) condition, and higher compared with the Heroin 1 mg/mL condition following Heroin 50 or 100 mg/mL (120-150 minutes) inhalation.

#### 3.1.3 Subcutaneous injection

##### Female Rats

The body temperature of the female rats (**Figure 4A**) was affected by Heroin injection, as confirmed by a significant effect of Time after injection [F (9, 54) = 15.21; P<0.0001] and of the interaction of Time with Heroin Dose [F (27, 162) = 2.40; P<0.0005] on temperature in the ANOVA. The post-hoc test confirmed that temperature was higher 120 minutes after injection of Heroin (0.5, 1.0 mg/kg), and lower 240-270 minutes after injection of 1.0 mg/kg, compared with the vehicle injection condition at the respective time. (N.b., we report only the key post-hoc results in text, additional findings for this entire experiment are depicted on the Figure panels.)

**Figure 4:**
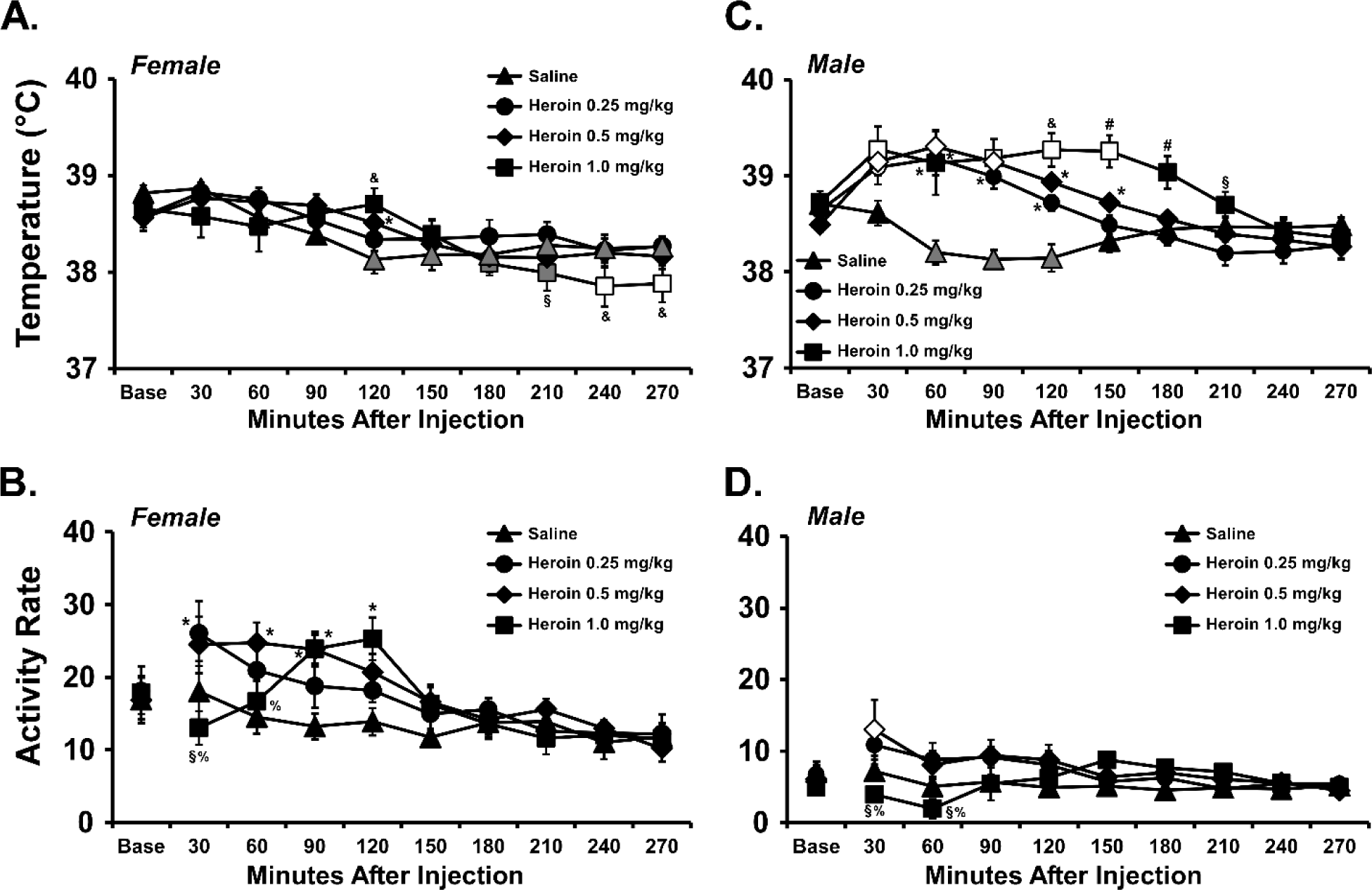
Mean (±SEM) body temperature (A, C) and activity rate (B, D) for female (N=7; A, B) and male (N=6; C,D) Sprague-Dawley rats after injection of saline or Heroin (0.25, 0.5, 1.0 mg/kg, s.c.). Shaded symbols indicate a significant difference from the baseline (Base) value, open symbols indicate a significant difference from the baseline and the Saline condition at the respective time-point. A significant difference from the Saline condition at a given time is indicated with *, a difference from Heroin 0.25 mg/kg with §, a difference from Saline and Heroin 0.25 mg/kg with &, a difference from Heroin 0.5 mg/kg with %, and a difference from all other treatment conditions with #.

The activity of the female rats (**Figure 4B**) was affected by Heroin injection, as confirmed by a significant effect of Time after injection [F (9, 54) = 14.01; P<0.0001], of Heroin Dose [F (3, 18) = 4.90; P<0.05] and of the interaction of Time with Heroin Dose [F (27, 162) = 2.36; P<0.001] on activity in the ANOVA. The post-hoc test confirmed that the activity was higher, in comparison with the respective time after vehicle injection, following 0.25 (30 minutes), 0.5 (60-90 minutes) or 1.0 (90-120 minutes) mg/kg. Activity was also significantly lower after the injection of 1.0 mg/kg compared with the injection of either 0.25 (30 minutes post-injection) or 0.5 mg/kg (30-60 minutes).

##### Male Rats

The body temperature of the male rats (**Figure 4C**) was significantly affected by Heroin injection,, as confirmed by a significant effect of Time after injection [F (9, 45) = 13.44; P<0.0001], of Heroin Dose [F (3, 15) = 16.13; P<0.0001] and of the interaction of Time with Heroin Dose [F (27, 135) = 7.14; P<0.0001] on temperature in the ANOVA. The post-hoc test confirmed that temperature was increased relative to baseline after injection of 0.25 mg/kg (30 minutes post-injection), 0.5 mg/kg (30-90 minutes), and 1.0 mg/kg (30, 90-150 minutes) and increased relative to the same time after saline injection by 0.25 mg/kg (30-120 minutes post-injection), 0.5 mg/kg (30-150 minutes), and 1.0 mg/kg (30-180 minutes).

The activity of the male rats (**Figure 4D**) was significantly affected by Heroin injection, as confirmed by a significant effect of Time after injection [F (9, 45) = 2.72; P<0.05] and of the interaction of Time with Heroin Dose [F (27, 135) = 2.32; P<0.001] on activity in the ANOVA. Activity was significantly increased 30 minutes after injection of 0.5 mg/kg relative to the baseline and the respective time after saline injection. Activity was also significantly lower 30-20 minutes after the injection of 1.0 mg/kg compared with the injection of either 0.25 or 0.5 mg/kg.

### 3.2 Experiment 2 (Anti-nociception)

When assessed after the exposure sessions, tail withdrawal latency in the male and female rats was significantly increased by heroin vapor inhalation compared with both the baseline assessment and after the PG inhalation condition (**Figure 5**). The effect was robust since the Heroin 50 and 100 mg/mL for 30-minute conditions resulted in maximal effects in different subsets of 5/7 male rats and 5/7 female rats each. That is, five animals in each sex failed to withdraw the tail within the 15 second time limit after each inhalation condition, but it was not the same five individuals across conditions. The 15-minute, Heroin 50 mg/mL condition resulted in a maximum effect in 1/7, and the 100 mg/mL for 15 minutes in 2/7, female rats.

**Figure 5:**
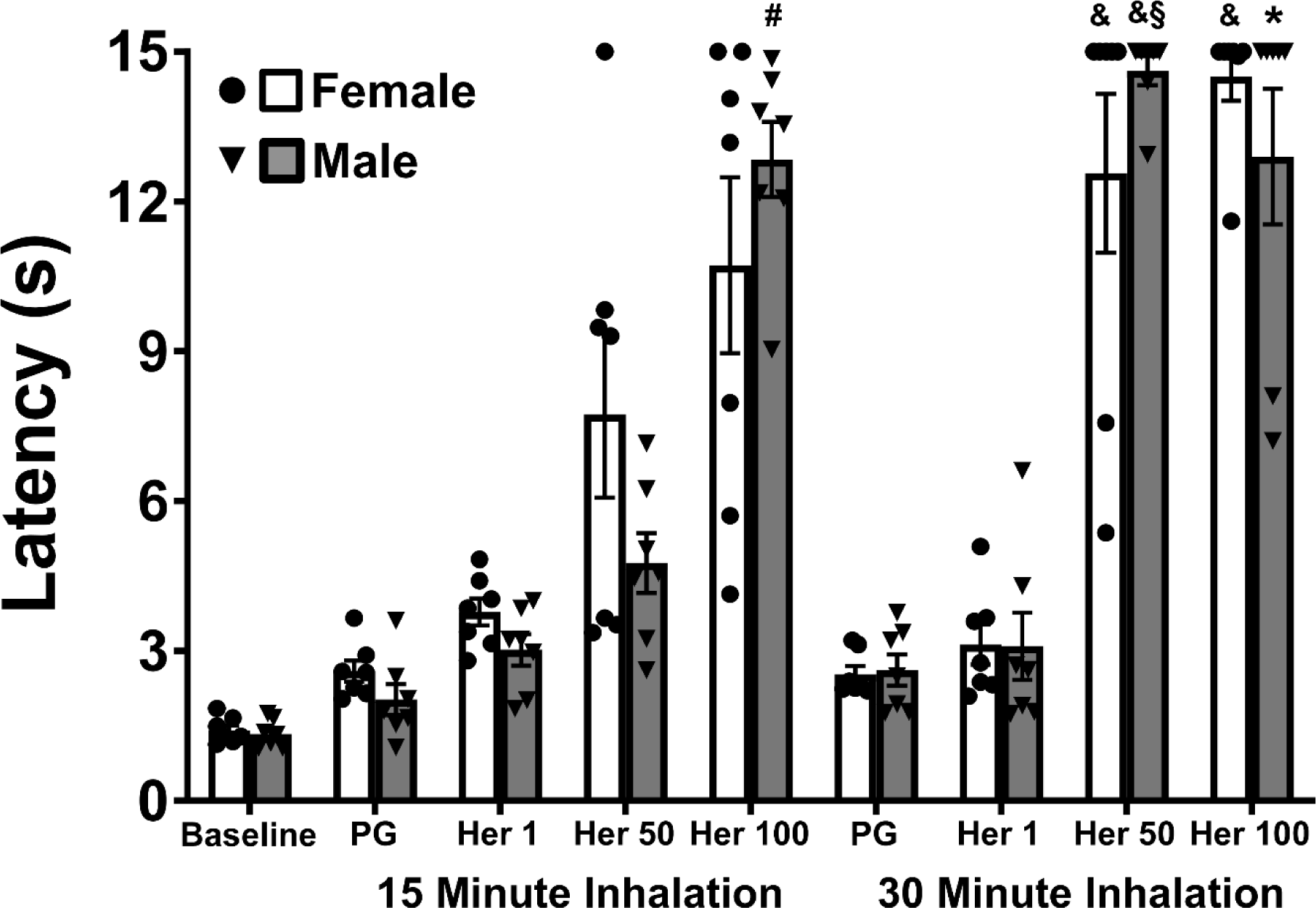
Mean (±SEM), and individual, tail withdrawal latency for male and female (N=7 per group) rats under baseline and post-vapor conditions. The baseline was conducted before any of the counter-balanced inhalation conditions and was not included in the statistical analysis. Within inhalation duration, a significant difference from the PG condition is indicated with *, a difference from PG and Heroin 1 mg/mL with &, and a difference from all other inhalation conditions with #. A difference between inhalation intervals within drug concentration is indicated with §. Base, baseline. PG, propylene glycol. Her 1, Heroin (1 mg/mL); Her 50, Heroin (50 mg/mL); Her 100, Heroin (100 mg/mL).

The three way ANOVA (excluding baseline) confirmed a significant effect of Vapor condition [F (1.857, 22.28) = 142.4; P<0.0001], of Inhalation Duration [F (1.000, 12.00) = 43.88; P<0.0001], of the interaction of Vapor Condition with Duration [F (2.157, 25.88) = 12.66; P<0.0005] and of the interaction of Sex, Vapor Condition and Duration [F (3, 36) = 3.33; P=0.0301], on tail withdrawal latency. The post-hoc test confirmed significant elevations relative to the respective PG and Heroin 1 mg/mL condition for the 50 and 100 mg/mL for 30-minute conditions in female rats and for the 50 mg/mL for 30 minutes conditions in male rats. There was also a significant difference in withdrawal latency in the male group following inhalation of Heroin 50 mg/mL for 30 minutes compared with the same concentration for 15 minutes. Tail withdrawal was also significantly slower in male rats after the inhalation of 100 mg/mL for 15 minutes compared with all other 15-minute conditions and slower after the inhalation of Heroin 100 mg/mL for 30 minutes compared with the PG for 30 minutes.

Follow-up analysis that was limited to the 15-minute conditions to further parse the three-way interaction failed to confirm any sex differences. Collapsed across sex, however, the Heroin 50 mg/mL condition significantly elevated tail-withdrawal latency compared with either the PG or 1 mg/mL condition. There was also a significant difference in withdrawal latency between the Heroin 50 mg/mL and each of the PG and Heroin 1 mg/mL conditions for the female rats.

## 4. Discussion

### 4.1 Main Findings

This study confirms that this method of inhalation of heroin, based on vapor created by an Electronic Drug Delivery System (EDDS) device, produces significant effects *in vivo* in both male and female rats. The effects are dose-dependent, as they vary with both inhalation duration (at a fixed concentration) and with drug concentration in the e-cigarette vehicle (at a given inhalation duration). We have previously shown that heroin vapor exposure (100 mg/mL, 30 minutes) produces moderate anti-nociceptive effects in male and female Wistar rats (Gutierrez et al., 2020) and that is extended to Sprague-Dawley rats, and to a wider range of inhalation conditions, in the present study. As we’ve previously shown, differences between common laboratory rat strains can result in quantitative or even qualitative differences in response to vapor inhalation of drugs, effects which may vary depending on the outcome measure (Taffe et al., 2020). The present results obtained near-maximal anti-nociceptive effects in more animals in the 100 mg/mL condition, compared with that prior study, however there were equipment differences between the two studies which may have resulted in more effective drug delivery. For example that prior study reported no anti-nociceptive effect of oxycodone vapor (100 mg/mL, 30 minutes) inhalation using first-generation Protank 3 canisters, whereas a study using the second-generation Herakles Sub Ohm canisters, more similar to the present second-generation SMOK canisters, did report anti-nociception subsequent to 100 mg/mL oxycodone vapor inhalation (Nguyen et al., 2019). Although vapor cloud density and chamber fill can be roughly equated by eye, it may be the case that different devices generate different droplet sizes or drug concentrations per droplet that produce complex interactions of a given drug with a given device to effect in vivo drug delivery. In that prior study a 1 mg/kg, s.c., heroin injection produced anti-nociception comparable to the effects of 30 minutes of 50-100 mg/mL exposure in this study. The data also show that the threshold for statistically reliable effects is 15-minutes of exposure to the 50 mg/mL condition. These exposure parameters led to significant anti-nociception (both collapsed across sex and within the female group) and produced significant effects on body temperature in the female rats. This study therefore establishes 50-100 mg/mL as an effective concentration range, and 1 mg/mL as an ineffective concentration, for 15-30 minutes of non-contingent inhalation exposure to heroin in rats.

This is a nontrivial advance from the prior demonstrations that rats and mice, respectively, will self-administer (or at least self-dose, this distinction is a longer discussion not directly relevant to the point at hand) the potent opioids sufentanil (Vendruscolo et al., 2018) and fentanyl (Moussawi et al., 2020). First, the doses needed to produce robust changes in nociception, thermoregulation and locomotor behavior are often far in excess of the doses that rodents will self-administer. Second, we found previously that cocaine and some amphetamine or cathinone class psychomotor stimulants which exhibit both low potency and lower solubility in PG may not readily be delivered in active doses with this approach (Nguyen et al., 2016a). Thus, it was critical to show that a less potent opioid such as heroin can be delivered with this method.

There were biphasic dose-dependent effects of inhaled heroin on body temperature which was expressed as lower exposure conditions (doses) producing reliable increases in temperature, and higher doses / exposures initially reducing body temperature, at least in the female rats. A prior study found similar biphasic effects after intravenous fentanyl, which initially decreased body temperature of male Long-Evans rats in the first hour after administration, but induced *hyper*thermia thereafter (Solis et al., 2017). This may reflect potency, or brain penetration speed, as potential differences between the two opioids that permitted this initial hypothermia to be observed. It could similarly be the case in the present study that the inhaled route of administration speeds the brain entry of heroin versus the more common s.c. or i.p. or even i.v. routes of administration. The observation that injected heroin did not have immediate effects on female rats’ body temperature (**Figure 4A**), and induced less complete suppression of activity (**Figure 4B**) compared with inhaled heroin (**Figure 2B**), supports this interpretation. If so, this may be a key area in which the development of this inhalation model allows the investigation of effects of heroin which are unique to this route of administration. It may also be the case that effects of inhalation are more similar to the effects of the intravenous route, in this case the method offers several practical advantages over implanting and maintaining intravenous catheters in rodents. Such advantages include overall cost, surgical expertise and avoiding subject loss due to occluded catheters or health complications related to the catheter.

The apparent sex difference (males did not express the initial hypothermia under any of the conditions tested) is most likely a difference in effective dose when heroin is inhaled, since pilot work with a group of male Sprague-Dawley rats illustrated a consistent *hypo*thermia associated with a high exposure condition and hyperthermia with a lower exposure condition (not shown). Also, female rats in this study were affected in terms of body temperature and anti-nociception after 15-minute inhalation of Heroin 50 mg/mL, whereas male rats were only affected on the anti-nociception assay and only to an extent that failed to reach statistical significance. Overall, these patterns are more consistent with a dosing difference across sex rather than a sex difference per se; additional work with more expanded dosing conditions might further explicate this interpretation. Interestingly, when mg/kg equivalent doses were injected the males’ body temperature was consistently elevated whereas the females’ temperature was not. A previous study found no difference in the ED50 for heroin (s.c.) in a warm water tail-withdrawal test between male and female Sprague-Dawley rats (Peckham and Traynor, 2006) and where sex differences were found (morphine, hydrocodone, hydromorphone) the males were *more* sensitive. Nevertheless, female mice develop greater hypothermia than do males after a mg/kg equivalent injection of morphine (Kest et al., 2000).

The initial suppression of activity, followed by a rebound of increased activity, is similar to that reported for injected morphine (Craft et al., 2006) but differs somewhat from prior results for injected heroin, which appeared to show a monotonic effect of time where activity is highest at the first time-points observed after dosing and then declined steadily with time. The pattern of initial suppression followed by a rebound was observed in the 1.0 mg/kg, s.c. injection study, as well as in the 30-minute inhalation experiments, suggesting the difference is not due to route of administration. One possible difference is that some of those studies were conducted with rats in the light part of the cycle (Hoffman and Wise, 1993; Swerdlow and Koob, 1985; Swerdlow et al., 1985), however others reporting similar monotonic time-courses were on a reverse cycle and were tested in the dark (Singh et al., 2005; Sorge and Stewart, 2006). A more consistent difference is that many studies of the effects of injected heroin used a non-housing, specialized photo cell cage (Hoffman and Wise, 1993; Singh et al., 2005; Sorge and Stewart, 2006; Swerdlow and Koob, 1985; Swerdlow et al., 1985) with mesh or rod floors, unlike the plastic floored normal housing cages with a thin layer of bedding used here. It may be the case that the more familiar, housing-type environment facilitated expression of a more naturalistic response.

There are a few necessary caveats; given the selected repeated measures experimental design it is always possible that a degree of plasticity, either sensitization or tolerance, may have developed. The counterbalanced testing order, however, minimizes the concern that this would have a systematic effect on any specific dosing conditions. Likewise, the animals had participated in prior studies involving exposure to THC, which might potentially produce cross tolerance, and because of that the animals were in the middle adult age range. It is entirely possible that responses in younger adults might differ. We have previously shown, however, that with ∼weekly gaps between successive THC exposure, male and female rats do not become tolerant to the hypothermic effects of THC and nor does the magnitude differ across 20-25 weeks of adult age (Javadi-Paydar et al., 2018a). In fact, it requires twice daily exposure to vapor inhalation for four days, at a minimum, to produce statistically significant acute tolerance (Nguyen et al., 2020b; Nguyen et al., 2018). Furthermore, a lasting-tolerance inducing regimen of THC exposure during adolescence does not alter the intravenous self-administration of oxycodone (Nguyen et al., 2020b) which again suggests that cross-tolerance/sensitization would not have resulted from the prior intermittent exposure of the animals to THC. It is likely that a similarly broad ranging set of studies would be required to parse the potential for intermittent acute heroin exposures similar to the present studies to induce plasticity of the response. For the present purposes of illustrating feasibility and dose control, however, such concerns are unlikely to affect the overall conclusions. Finally, opioids such as heroin suppress respiration at high doses, which might complicate the uptake of drug in an inhalation paradigm. This is unlikely to be the case for the present studies since a 7.5 mg/kg intravenous dose, or a 10 mg/kg intraperitoneal dose, fails to alter tidal volume in mice (Hill et al., 2020). A 90 mg/kg, i.p. dose was required to reduce tidal volume in the Hill et al. (2020) study and a toxicological report shows that 21.5 mg/kg, i.p. heroin is lethal to only 67% of rats (Strandberg et al., 2006). Animals in this study were not in clinical respiratory distress observable to expert research staff on removal from the inhalation chambers. Combined, this evidence suggests that any ongoing respiratory effects of heroin were unlikely to be large enough to alter drug uptake across the session.

### 4.2 Alternate Methods

This study reports a set of methods we have evolved from our ongoing experience using EDDS-based vapor exposure. Through the developmental arc of this approach, it is clear that this is but one specific set of methods and that many variations are likely to also work in the hands of other interested laboratories. First, we have used several different iterations of e-cigarette canister/atomizer and found that most differences are subtle in terms of drug delivery. There is a constantly changing and highly varied array of products available online and in local stores and most of them are likely to function for laboratory purposes. Effective vapor delivery can be achieved with many different types of cannister / atomizer products, with only modest methodological adjustments of puffing schedule, drug concentration and inhalation time.Second, the volume of the sealed chamber used to expose animals and the number per chamber can be varied, depending on the goals to be attained and the available equipment. The initial validation step is simply to determine that a vapor cloud to fill the available chamber volume can be produced. Third, while we have adopted a puffing schedule of every 5 minutes, more-, or less-frequent puffing could be used. The most important consideration for developing these methods is to have the ability to contrast and validate the effects of manipulating drug exposure parameters. Validation should be done with a desired target endpoint (whether that be anti-nociceptive effects, effects on spontaneous behavior, on physiological endpoints such as thermoregulation, or other measure of interest) that is reliably produced in the relevant laboratory species/strain/genotype by drug injection, or other route of drug administration. Once the target endpoint is determined, systematic manipulation of equipment, dosing protocols or drug concentrations can be used to achieve dose control.

## 5. Conclusions

In conclusion, the e-cigarette based vapor exposure approach delivers physiologically and behaviorally relevant amounts of heroin to rats. Given the growing opioid crisis, and the widespread use of e-cigarette technologies, it will be of increasing utility to have a rodent pre-clinical model for the assessment of the effects of inhaled heroin and other opioids.

## Acknowledgements

The study was conducted with the support of United States Public Health Service research grants from the National Institutes for Health (R01 DA035281, R01 DA042211), by aa T32 AA007456 (Gutierrez, Fellow), by a UCSD Chancellor’s Post-doctoral Fellowship Program (Gutierrez, Fellow), and the Tobacco-Related Disease Research program of California (T31IP1832). The NIH/NIDA and the TRDRP had no role in study design, collection, analysis and interpretation of data, in the writing of the report, or in the decision to submit the paper for publication. The authors declare no additional financial conflicts which affected the conduct of this work.

## Declarations of interest

none.

## Notes

### Competing Interest Statement

The authors have declared no competing interest.

### Summary of Updates

Revisions provided: -expand upon a few methodological details -add an additional data set from heroin injections for comparison with the inhalation -provide individual points for the nociception figure -a new caveats paragraph in the Discussion

